# Phylogenomic Insights into the Evolution and Origin of Nematoda

**DOI:** 10.1101/2023.12.13.571554

**Authors:** Xue Qing, Y. Miles Zhang, Sidi Sun, Mohammed Ahmed, Wen-Sui Lo, Wim Bert, Oleksandr Holovachov, Hongmei Li

## Abstract

The phylum Nematoda represents one of the most cosmopolitan and abundant metazoan groups on Earth. In this study, we reconstructed the phylogenomic tree for phylum Nematoda. A total of 60 genomes, belonging to eight nematode orders, were newly sequenced, providing the first low-coverage genomes for the orders Dorylaimida, Mononchida, Monhysterida, Chromadorida, Triplonchida, and Enoplida. The resulting phylogeny is well-resolved across most clades, with topologies remaining consistent across various reconstruction parameters. The subclass Enoplia is placed as a sister group to the rest of Nematoda, agrees with previous published phylogenies. While the order Triplonchida is monophyletic, it is not well-supported, and the order Enoplida is paraphyletic. Taxa possessing a stomatostylet form a monophyletic group; however, the superfamily Aphelenchoidea does not constitute a monophyletic clade. The genera *Trichinella* and *Trichuris* are inferred to have shared a common ancestor approximately 202 mya, a considerably later period than previously suggested. All stomatostylet-bearing nematodes are proposed to have originated ∼305 mya, corresponding to the transition from the Devonian to the Permian period. The genus *Thornia* is placed outside of Dorylaimina and Nygolaimina, disagreeing with its position in previous studies. Additionally, we tested the whole genome amplification method and demonstrated that it is a promising strategy for obtaining sufficient DNA for phylogenomic studies of microscopic eukaryotes. This study significantly expanded the current nematode genome dataset, and the well-resolved phylogeny enhances our understanding of the evolution of Nematoda.

## Introduction

The phylum Nematoda represents one of the most cosmopolitan, abundant, and diverse metazoans on Earth, with an estimated number of species extending to 10 million (Lambshead, 1993) (Figure 1). Free-living nematodes occupy all trophic levels in the food web and play a pivotal role in carbon cycling and energy flows (Ferris, 2010; Ingham et al., 1985; van den Hoogen et al., 2019), and the genera *Caenorhabditis* and *Pristionchus* are two popular nematode models in molecular and evolutionary biology. Nematode parasites are a threat to not only the health of plants (e.g. *Meloidogyne* spp., *Heterodera* spp.), but also that of animals and humans (*e*.*g*. ascariasis, trichuriasis, and hookworm disease, filariasis). Resolving nematode phylogeny is an important step in understanding the origin of parasitism and transition of lifestyles, from basic evolutionary biology to pathogen control and development of anthelmintic drugs (International Helminth Genomes Consortium, 2019). However, the high phenotypic plasticity (Coomans, 2002), a near-complete lack of synapomorphic diagnostic characters, and the absence of an informative fossil record have prevented scientists from deriving a consistent evolutionary framework (Lorenzen, 1981; Coomans, 2000). The first molecular phylogeny was reconstructed based on 18S rRNA of 53 species and led to the recognition of three major lineages of nematodes: Dorylaimia, Enoplia, and Chromadoria (Blaxter et al.,1998; De Ley and Blaxter et al., 2002). However, despite its popularity as a molecular marker, 18S rRNA lacks the resolving power for deep phylogenies, i.e. the root of the nematode tree (Blaxter and Koutsovoulos, 2015). As a result, currently all published rRNA phylogenies lack supporting evidence for the “backbone” of the tree, and many lineages remain poorly resolved (Ahmed and Holovachov, 2021).

**Figure 1.**
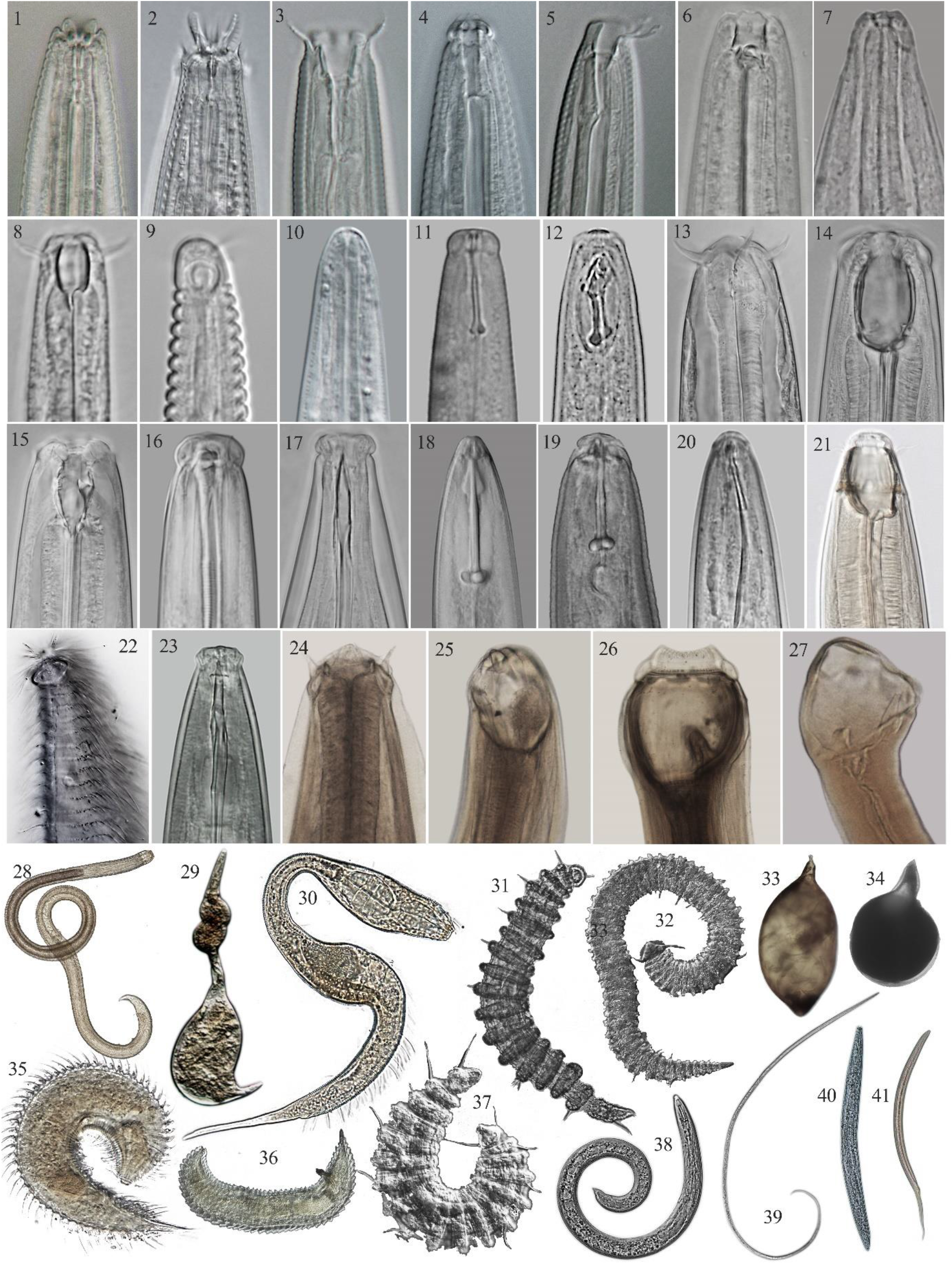
Diversity of free-living and parasitic species in Nematoda. 1-27: the head region, 28-35: body habitus. 1, 40:*Acrobeloides*; 2: *Acrobeles*; 3: *Bicirronema*, 4: *Teratocephalus*; 5: *Tricirronema*; 6: *Pristionchus*; 7: *Steinernema*; 8: *Prismatolaimus*, 9: *Aphanolaimus*, 10: *Alaimus*; 11: *Aphelenchoides*; 12: *Diphtherophora*; 13: *Tripyla*; 14: *Miconchus*; 15: *Mylonchulus*; 16: *Ironus*; 17: *Aporcella*; 18, 38: *Helicotylenchus*; 19, 34: *Meloidogyne*; 20: *Trichodorus*; 21: *Sphaerolaimus*; 22, 35: *Greeffiella*; 23: *Eudorylaimus*; 24: *Oesophagostomum*; 25: *Ancylostoma*; 26: *Strongylus*; 27: *Bunostomum*; 28: *Croconema*; 29: *Tylenchulus*; 30: Draconematidae; 31, 37: *Desmoscolex*; 32, 36: *Tricoma*; 33: *Heterodera*; 39: *Etamphidelus*; 41: *Malenchus*. 7: insect parasite; 18-20, 29, 33, 34, 38: plant parasites; 24-27: vertebrate parasites; 1-6, 8-17, 23, 39-41: terrestrial free-living species; 21, 22, 28, 30-32, 35-37 marine free-living species.

By contrast, phylogenomic analyses have yielded statistically-robust and congruent results that have proven useful in resolving previously controversial phylogenetic relationships across evolutionary lineages (Misof et al., 2014; Telford et al., 2015; Williams et al., 2020). Advances in sequencing technology have also enabled the generation of inexpensive low-coverage genomes for phylogenomic studies (Zhang et al. 2019, Kim et al. 2024). However, difficulties in obtaining high-quality DNA have remained a major challenge for -omics studies on uncultured microscopic organisms. In the case of nematodes, pioneering work has been limited to studies on vertebrate parasitic taxa and model *Caenorhabditis* spp. (Blaxter and Koutsovoulos, 2015; International Helminth Genomes Consortium, 2019; Stevens et al., 2019), with little or no representation for species less than 1mm in length as well as for uncultured species. Recent advances in whole-genome amplification (WGA) have enabled single cells to generate sufficient DNA for sequencing, and these techniques have now been applied to various microscopic eukaryotic (Lepere et al.2011; Weisz et al., 2019; Sahraei et al., 2022; Lee et al., 2023), making it possible to sequence nematodes from a single individual at a phylum scale. The current paper focuses on sequencing the low-coverage genomes of 60 nematode species covering eight orders using WGA, aiming to provide finer resolution for phylogeny and improved knowledge about patterns and processes of nematode evolution.

## Material & Methods

### DNA extraction, sequencing, and genome assembly

Nematodes were isolated from soil or marine sediments and extracted using a Baermann funnel. One to five adult fresh worms were mounted on temporary slide and morphologically identified under an Olympus BX51 microscope (Olympus Optical, Japan). The coverslip was then removed, the worms were cut into pieces with a sterilized scalpel, and transferred with a needle into a 200 μl PCR tube. Genomic DNA extraction and amplification were performed with the REPLI-g Single Cell Kit (Qiagen, Germany) using 1–12 individual alive nematodes. In addition to the whole genome amplification (WGA) method, DNA was directly extracted from 20,000 fresh individuals for three species (*Aphelenchoides blastopthorus, Aphelenchoides smolae* and *Ditylenchus* sp.) using the Ezup Column Animal Genomic DNA Purification Kit (Sangon Biotech, China), in order to evaluate the performance of WGA. The TruSeq DNA Sample Prep Kit (Illumina Inc. CA, USA) was used for library preparation according to the manufacturer’s instructions. Sequencing was preformed using the Illumina NovaSeq platform. Quality control was performed in FASTP (Chen et al., 2018), reads less than 50 bp, Q-score less than 20, or reads with N base number > 3 were removed. Filtered reads were assembled using SPAdes v3.15.5 with the default parameters (Bankevich et al. 2012). We ran BlobToolKit (Challis et al., 2020) on raw assemblies to manually screen for scaffolds derived from non-target organisms. To avoid potential noise, we removed fragments with total length < 3k and coverage < 15. The remaining sequences were subjected to a microbial contamination prediction analysis using Kraken2 (Wood et al., 2019) to identify and eliminate any fragments stemming from microbial contaminants. The raw sequences generated in this study were submitted to GenBank SRA database with accession number PRJNA1128552.

### Phylogenomic analysis

The phylogenetic hypothesis was reconstructed using newly generated data supplemented with additional genomes and transcriptomes downloaded from GenBank (see Table S1), and a total of 156 Nematoda and ten outgroups (two of each from Phyla Nematomorpha, Tardigrada, Arthropoda, Onychophora, and Priapulida) were included. The complete and single copy ortholog genes were extracted using BUSCO (Simão et al. 2015) using the Nematode ortholog database OrthoDB v10 2024-01-08, and translated into amino acids (AA). We selected a 50% (USCO50, 1365 loci) and 70% complete (USCO70, 191 loci) matrices using custom script *matrix_generation*.*sh* from Du et al. (2023) for downstream analyses to balance between the number of loci and data completeness. The matrices were aligned using the L-INS-I option in MAFFT v7.490 (Katoh and Toh, 2008), trimmed using TrimAl (Capella-Gutiérrez, et al. 2009), and concatenated using FASconCAT-G v1.05.1 (Kück, 2014). We further removed potentially misaligned regions using SpruceUp (Borowiec, 2019) at 97% lognormal criterion. The remaining loci were further filtered using custom script *loci_filtering_alignment-based*.*sh* provided by Du et al. (2023) wrapper for PhyKIT v1.11.10 (Steenwyk et al., 2021) to remove loci with <100% parsimony-informative sites (Shen et al., 2016); loci with <0.35 relative composition variability (Phillips and Penny, 2003); and loci failing symmetry test (*p*-value 0.01-0.1) for stationarity and homogeneity (Naser-Khdour et al., 2019). The summary statistics of the final filtered matrices were calculated using AMAS v0.98 (Borowiec, 2016).

The following analyses were performed on both matrices using IQ-TREE v2.2.0 (Minh et al., 2020a): 1) Partitioned by locus with best model selected using ModelFinder (Kalyaanamoorthy et al. 2017), and with ultrafast bootstrap (Hoang et al. 2017) and SH-like approximate likelihood ratio test (SH-aLRT, Guindon et al. 2010) as support values; 2) Individual gene trees with best model selected using ModelFinder; 3) Gene and site concordance factors (gCF, sCF, respectively) using steps 1 and 2 following (Minh et al. 2020b); 4) GHOST model to account for heterotachy (Crotty et al. 2019); and finally, 5) the posterior mean site frequency model (PMSF) which has been designed to correct for long-branch-attraction by modeling site heterogeneity (Wang et al. 2018), the C20 model was only used for USCO50 matrix due to the computational limits required for such a large dataset, whereas the more complex C60 model was used for the USCO70 matrix.

We also analysed both datasets using the multispecies coalescent-based method (MSC) implemented in ASTRAL-III (Zhang et al. 2018), using gene trees generated in step 2 above as input. We collapsed the gene tree nodes with <10% support using Newick Utilities (Junier, 2010), and support was evaluated using local posterior probability (LPP, Sayyari and Mirarab, 2016). Detailed scripts and flags for both IQ-TREE and ASTRAL can be found in Table S2.

### Molecular dating and ancestral state reconstruction

Divergence dating was performed using MCMCTree using PAML v4.9j (Yang, 2007) using a subset of the PMSF_C20 topology as guide tree and the USCO50 matrix, filtered using the custom script *loci_filtering_tree-based*.*sh* provided by Du et al. (2023) based on the gene trees built in step 2 above with the following: average bootstrap support of >80 (Salichos and Rokas, 2013); signal-to-noise ratio (treeness/relative composition variability) of 1.5 (Phillips and Penny, 2003); and degree of violation of the molecular clock of 0.2 (Liu et al., 2017); to obtain a final subset matrix of 135 loci. Approximate likelihood calculation and ML estimation of branch lengths to reduce the computational burden by calculating the Hessian matrices using the LG substitution model and the independent rates clock mode. Given the controversial relationships among Cryptovermes (Nematoida and Panarthropoda) within Ecdysozoa is beyond the scope of this study, we chose not to include fossil outgroups from these lineages. Instead we used fossil age from Phylum Priapulida dated at 518 mya as a calibration point for Scalidophora which is the sister to Cryptovermes, and a root age of 636 mya following Howard et al. (2022). Seven nematode fossils plus one Priapulida fossil were selected for calibration based on existing fossil records (Table S3).

Two independent runs were performed with 50,000 generation burnins and 500,000 generations using the HKY85 model, and convergence was assessed using Tracer v1.7 (Rambaut et al. 2018). The tree was visualized using MCMCtreeR (Puttick, 2019) in R v4.2.0 (R Core Team 2022).

## Results

### Whole genome amplification and genome assembly

A total of 60 species from terrestrial and marine environments were newly sequenced. The raw assemblies ranged in size from 43.67 Mb to 839.99 Mb, comprising contigs spanning from 15.99 Kb to 2.69 Mb. Genome sizes decreased after removing potential contaminations, with lengths ranging from 2.90 MB to 189.63 Mb, and contig counts varying from 484 to 33537 (Table S4). The BUSCO analysis yielded completeness percentages ranging from 7.65% to 81.86% for the assembled genomes. We observed smaller size and more fragmented genomes (higher contig numbers) when using the WGA method compared to direct DNA extraction without amplification from fresh nematodes (Figures S1, S2). Further BUSCO evaluation indicated varying performance results, *i*.*e*. both methods generated a similar genome completeness in *Ditylenchus* sp., a more complete *A. blastopthorus* and *A. smolae* genome with direct extraction and WGA, respectively (Figure S3). The assembled contigs were annotated and plotted according to their GC content and coverage (Figure S4), revealing that direct extraction introduced more bacterial contaminations than the WGA approach.

### Evolution of Nematoda

From the assembled genomes, we extracted 1028 AA loci with 304,894 bp for the USCO50 matrix, and 115 loci with 23,069 bp for the USCO70 matrix. In general, the topologies inferred from USCO70 matrix were less supported when compared to those from the USCO50 matrix. The reconstructed phylogenies have similar topologies in most cases regardless of the metrics, datasets, and reconstruction methods (Figure 2). Nematoda is either sister to Nematomorpha (PMSF) or to a clade including Nematomorpha and Tardigrada (MSC and GHOST). Within Nematoda, the differences between topologies are summarised in the table of Figure 2. The subclass Enoplia primarily consists of taxa inhabiting marine environments. Placing the class Enoplia as the sister taxon to the rest of Nematoda in our analysis suggests a likely marine origin for the nematode ancestor. This subclass comprises two orders, Enoplida and Triplonchida. Notably, the representation of taxa in the latter group has significantly improved compared to transcriptomic analyses (Smythe et al. 2019; Ahmed et al. 2022). The monophyly of Triplonchida is well-supported. Enoplida is paraphyletic due to *Ironus dentifurcatus* being part of a lineage that is sister to the rest of Enoplida. This is consistent with the previous finding of Ahmed et al. (2022) who recovered Enoplida as paraphyletic, with *Bathylaimus* sp. (Triplonchida) and *Tobrilus* sp. (Enoplida) forming a sister branch to the rest of Enoplida.

**Figure 2.**
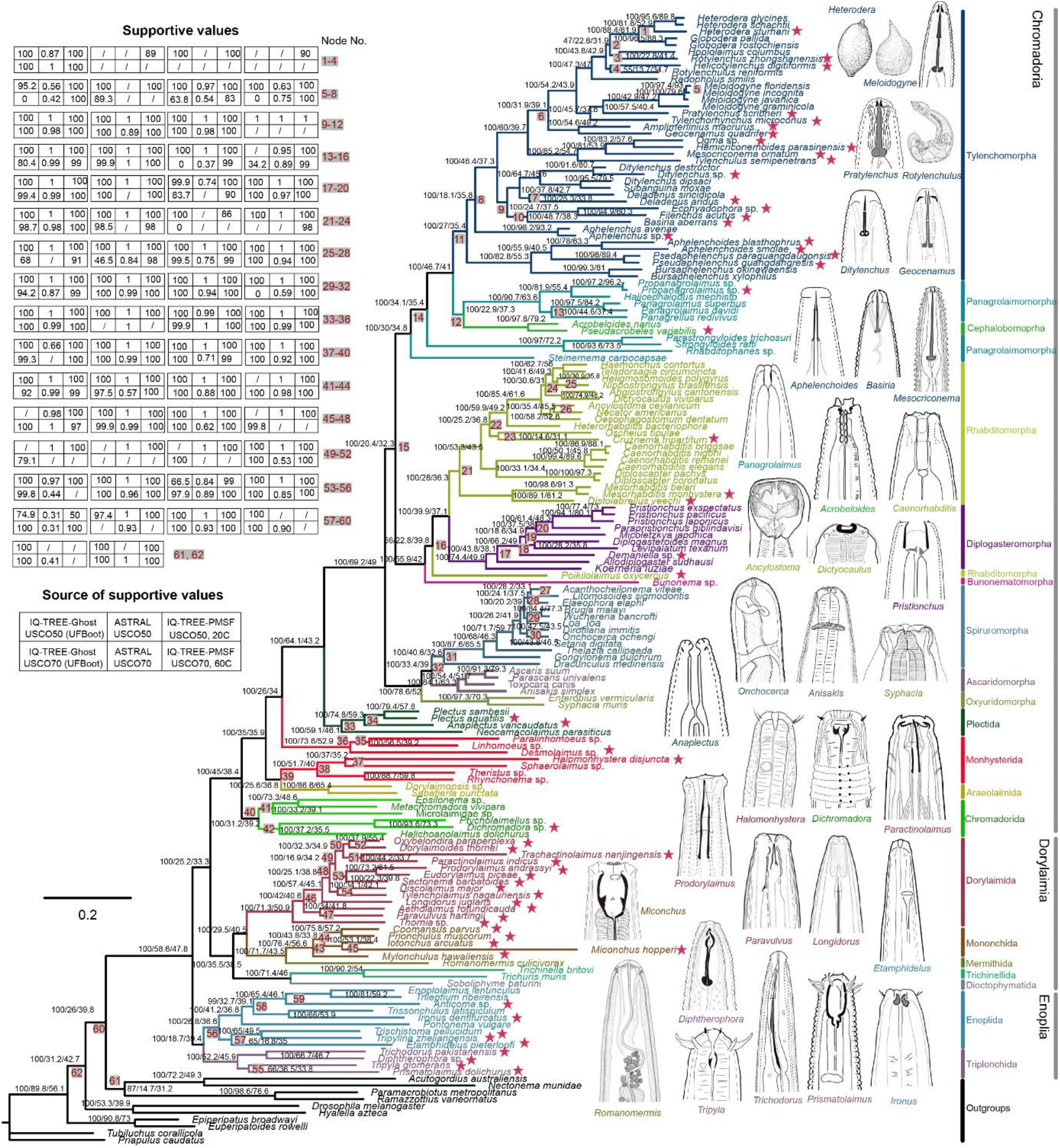
Phylogeny of Nematoda inferred from amino acids matrix USCO50 implemented in IQ-TREE. Node labels show ultrafast bootstrap approximation (UFBoot)/gene concordance factor (gCFs)/site concordance factor (sCFs). Node supports from other analyses are given in the table at left top corner, with the corresponding node numbers indicated by the coloured squares at tree node. The slash in table suggests the topology is not supported. The nodes without corresponding nodes number suggest they were fully supported in all analyses. The newly sequenced species were indicated with star.

There are five orders within Dorylaimia (clade I in Blaxter et al.,1998) represented in this analysis, all of which are terrestrial. A lineage consisting of the vertebrate parasitic orders Trichinellida and Dioctophymatida forms a sister branch with a clade comprising Dorylaimida, Mononchida and Mermithida. The orders Dorylaimida, Mononchida and Trichinellida, which are represented by more than one taxon, all form well-supported monophyletic groups, consistent with previous 18S rRNA-based and phylogenomic analyses (van Megen et al. 2009; Smythe et al. 2019; Ahmed et al. 2022). Apart from well-represented Rhabditida, four early-diverging Chromadoria orders are represented in this study, Chromadorida, Araeolaimida, Monhysterida and Plectida, with the first three being mostly marine. Within Rhabditida, all three suborders are strongly supported. The three infraorders representing Spirurina also received robust support. The Diplogasteromorpha within Rhabditina was likewise well-supported, but Rhabditomorpha was paraphyletic with genus *Poikilolaimus* being sister lineage to the clade of Diplogasteromorpha + Rhabditomorpha. The Bunonematomorpha was only represented by one genus *Bunonema*, and was placed either as sister taxa to all other members of Rhabditina (most cases) or to *Poikilolaimus* (GHOST model in IQ-TREE). Even though the analysis accounted for all three infraorders of Tylenchina, the majority of the taxa belonged to the infraorder Tylenchomorpha. In congruence with previous 18S rRNA and transcriptomic analyses of Tylenchina, Strongyloidoidea, one of the superfamilies of Panagrolaimomorpha, diverged first from the rest of Tylenchina, with two separate branching events–first with Steinernematidae, and then the Alloionematidae + Strongyloididae lineage. The remaining crown members of the tree formed a well-supported branch, within which Panagrolaimidae, another family of Panagrolaimomorpha, nested. Consistent with other phylogenomic studies, but contradictory to 18S rRNA data, Tylenchomorpha was found to be monophyletic, supporting the single evolutionary origin of the stomatostylet (the tylenchid and aphelenchid feeding apparatus), although the superfamily Aphelenchoidea does not form a monophyletic clade within Tylenchomorpha.

### Molecular dating of Nematoda

The divergence time dating, based on the ten calibrated nodes (Figure S5), traces the divergence date for Nematoda from its sister group Nematomorpha during late Ediacaran to Ordovician period (620–455 mya) (Table 1). Within Nematoda, the most recent common ancestor of all Enoplia dated back to a period spanning the end of the Ediacaran to the mid-Devonian (573–378 mya), slightly later than the common ancestor of all Chromadoria (561– 408 mya) in mean highest posterior distribution (HPD, 473 *vs* 482 mya). All Dorylaimia had their most recent common ancestor ∼446 mya, during the beginning Cambrian through to the end of Devonian. Our analysis indicates that human parasite *Trichinella* and *Trichuris* shared a common ancestor ∼202 mya, which is a considerably more recent compared ∼275 mya and ∼281 mya, respectively reported in Zarlenga et al. (2006) and Korhonen et al. (2016). The most recent common ancestor of all stomatostylet-bearing nematodes is suggested to have existed ∼305 mya, coinciding with the transition from the Devonian to end of the Permian period.

**Table 1.** The Nematoda divergence time estimation performed by using MCMCTree on the amino acids matrix USCO50 PMSF C20 topology.

| Clade | Time estimations (mya) |  |  |
| --- | --- | --- | --- |
|  | Mean HPD | 95% Lower HPD | 95% Upper HPD |
| Nematoda | 531 | 455 | 620 |
| Chromadoria | 482 | 408 | 561 |
| Spirurina (Clade III) | 323 | 259 | 388 |
| Spiruromorpha | 261 | 198 | 322 |
| Ascaridomorpha | 228 | 168 | 295 |
| Oxyuridomorpha | 148 | 66 | 224 |
| Tylenchina (Clade IV) | 361 | 302 | 427 |
| Cephalobomorpha | 105 | 45 | 178 |
| Tylenchomorpha | 305 | 254 | 364 |
| Rhabditina (Clade V) | 336 | 277 | 403 |
| Diplogasteromorpha | 240 | 185 | 296 |
| Strongyloidea | 120 | 90 | 154 |
| Plectida | 294 | 181 | 405 |
| Linhomoeina | 291 | 193 | 399 |
| Monhysterina | 325 | 238 | 413 |
| Chromadorida | 348 | 242 | 454 |
| Desmodorida | 345 | 238 | 456 |
| Dorylaimia (Clade I) | 446 | 367 | 537 |
| Dorylaimida | 290 | 219 | 363 |
| Mononchida | 210 | 148 | 278 |
| Trichinellida | 202 | 120 | 292 |
| Enoplia (Clade II) | 473 | 378 | 573 |
| Enoplida | 403 | 305 | 501 |
| Triplonchida | 304 | 194 | 420 |

## Discussion

### Evolution of early-branching Nematoda

The general topology of the tree agrees well with the results of previously-published analyses obtained using mostly transcriptomic datasets for different sets of species (Smythe et al., 2019, Ahmed et al., 2022), with Enoplia (Clade II of Blaxter et al., 1998) diverging first from the rest of the nematodes, and Dorylaimia (Clade I of Blaxter et al., 1998) being close second, in full agreement with the scarce morphological and embryological evidence summarised by Holterman et al. (2006). Within Enoplia, all included species are represented by newly sequenced genomes, and although none of the taxa are identical to earlier datasets, the resultant topology agrees in basic details with the transcriptome-only phylogenies (Smythe et al., 2019, Ahmed et al., 2022). However, the bootstrap support of several nodes within Enoplida is suboptimal: Triplonchida is resolved as monophyletic, while Enoplida as paraphyletic. This agrees with morphological evidence for the monophyly of Triplonchida suggested by De Ley and Blaxter (2002) but contradicts the phylogenies based on 18S rRNA gene (summarised in Ahmed and Holovachov, 2021). Enoplia (Clade II) is one of the most morphologically diverse groups of nematodes, so there are no easily testable morphology-based phylogenies. Enoplia are also likely to comprise the first nematode lineages that made the transition from marine to freshwater habitats in the Ediacaran-Devonian periods several times independently (Table 1). Future sampling of additional genomes from all lineages within Enoplia will not only help resolve relationships within the group but will also shed light on how colonisation of freshwater habitats took place in the early Palaeozoic.

The tree topology within Dorylaimia (Clade I) not only agrees fully with previously published datasets (Smythe et al., 2019, Ahmed et al., 2022), but a considerably expanded dataset of Mononchida and especially Dorylaimida also allows further comparison with morphology- and rRNA-based phylogenies. Within Dorylaimida, the genus *Thornia* branches off first, which disagrees with morphological evidence (Peña-Santiago, 2014). The sole available 18S rRNA sequence of *Thornia* is highly divergent and its position is therefore unreliable (Fig. S1. in Holterman et al. 2019). The remaining phylogeny of Dorylaimida is more in agreement with previous studies: Nygolaimina (represented here by *Paravulvus* and *Aetholaimus*) and Dorylaimina (all other taxa) are sister clades. The two families represented by more than one species are Actinolaimidae (*Paractinolaimus* and *Trachactinolaimus*) and Qudsianematidae (*Eudorylaimus* and *Discolaimus*), are herein resolved as monophyletic and polyphyletic respectively, in agreement with previous rRNA-based analyses (summarised in Ahmed and Holovachov, 2021). The entire Clade I is represented by limno-terrestrial (Mononchida and Dorylaimidae) and parasitic (Trichinellida, Dioctophymatida and Mermithida) species. Of the latter, only Trichinellida includes considerable number of parasites inhabiting marine vertebrate species. Molecular dating suggests that the Clade I diverged off from the rest of Nematoda in the Cambrian-Devonian, but the phylogeny of the group is not detailed enough to hypothesize about the transition between marine, freshwater, and terrestrial habitats within this lineage. Mermithida, parasitoids of insects and other (mostly) terrestrial arthropods are represented by single species in the current analysis; molecular dating suggests a very broad timeframe for the origin of the clade, spanning from Devonian to Triassic, much later than the period in which their arthropod hosts have diversified on land. Similarly, the plant-parasitic lineage within the Clade I is represented by a single species, *Longidorus juglans*, and molecular dating suggests it split off from non-parasitic dorylaims during the Permian-Jurassic, at about the same time as their hosts, flowering plants. However, more data on the evolution of both hosts and parasites is necessary to make in-depth cophylogenetic study.

### Evolution of plant-parasitic Tylenchina

Taxa closely related to plant-parasitic lineages have long been a focal point of interest in understanding the origins of plant parasitism (Bert et al., 2008; Ahmed et al., 2022). Most rRNA-based phylogenies indicate that Cephalobomorpha shares the most recent common ancestry with Tylenchomorpha (Blaxter et al., 1998; Holterman et al., 2006; Bert et al., 2008; van Megen et al., 2009). However, our analyses are consistent with the analyses of Ahmed et al. (2022), which positions both Cephalobomorpha and Panagrolaimomorpha as sister taxa to Tylenchomorpha. The taxonomic position of aphelenchs (Aphelenchoidea: Aphelenchoididae + Aphelenchidae) relative to the tylenchs (Tylenchomorpha without aphelenchs) has been a subject of especially extensive debate. Despite the evident morphological similarities between Aphelenchidae and Aphelenchoididae, the latter has been placed outside the Tylenchomorpha based on rRNA-based phylogenies. They were separated from Aphelenchidae, which were found to be sister to the remaining members of Tylenchomorpha (van Megen et al., 2009). This placement was hypothesized as resulting from long branch attraction and/or elevated AT-content (De Ley and Blaxter, 2002; Holterman et al., 2006) and their monophyletic grouping could not be significantly rejected (Bert et al., 2008). In contrast, mitogenome trees have consistently shown aphelenchs as monophyletic and positioned away from other Tylenchomorpha (Kern et al., 2020). This analysis supports the hypothesis that is most plausible based on morphology, namely that taxa equipped with the stomatostylet form a monophyletic group, a supposition that is congruent with the findings of Ahmed et al. (2022). Nonetheless, in our analysis aphelenchs are still depicted as paraphyletic and Aphelenchidae are, similar to rRNA-based phylogenies, more closely related to tylenchs than to the morphologically more similar Aphelenchoidea.

### Limitations of molecular dating of nematode phylogeny

Although the number of known nematode fossils is much larger than for some other soft-bodied invertebrate phyla (Poinar et al., 2011), their usefulness for molecular dating is limited. The large majority of known nematode fossils originate from various fossilised resins (amber and copal), are relatively recent and, due to the nature of the fossilisation, are strongly biased towards insect-associated and terrestrial nematodes. The next most common category also includes relatively recent fossils of vertebrate parasites found in coprolites and mummified remains. Only a few fossils date back to Palaeozoic, during which the majority of nematode lineages are thought to have diverged. Comparing the dates of origin of parasitic nematodes and their respective hosts we find both agreements and disagreements. The possible date of origin of Longidoridae, which is parasite of flowering plants, matches that of their hosts during the Jurassic-Cretaceous period, while plant-parasitic Tylenchomorpha has an earlier origin (Devonian to the Permian) than its hosts. The earliest possible divergence date of plant-parasitic Triplonchida, represented in our analysis by only *Trichodorus pakistanensis* (Carboniferous to early Cretaceous) predates the origin of their hosts, flowering plants (van der Kooi and Ollerton, 2020). Similarly, the possible date of divergence of animal parasitic Ascaridomorpha is underestimated in our analysis (early Permian to Jurassic) compared to cophylogenetic study using fossil data from their vertebrate hosts (Carboniferous) (Li et al., 2018). The dating of the split between *Trichinella* and *Trichuris* is also inconsistent between our and previous analyses (Zarlenga et al, 2006; Korhonen et al. 2016). These examples not only expose the limitations of the fossils when dating the nematode phylogeny, but highlight the need for a multidisciplinary approach, combining fossil evidence, co-phylogenetic studies, and mutation accumulation rates.

### Whole genome amplification and microscopic eukaryotes phylogenomic

Microscopic eukaryotes, such as Protozoa, Nematoda, Platyhelminthes, and Gastrotricha, are abundant, diverse and fill critical ecological roles across every ecosystem on Earth, and yet they were poorly known across subjects ranging from biodiversity and, evolution, to genomics (Wardle, 2006; Bik et al., 2012). The lack of omics studies in these groups can partly be explained by their small body size and low DNA quantity. Indeed, the high-throughput sequencing protocols require large amounts of input DNA (*e*.*g*. several micrograms), typically thousands of times more than that which is contained within a single microscopic individual. Zhang et al. (2019) proposed WGA in genome sequencing and phylogenomic of small animals, and three pioneering studies have used this method based on Illumina and PacBio platform to generate nematodes genomes (Roberts et al., 2024; Lee et al., 2023; Doyle et al., 2019), although no study had yet evaluated its performance using microscopic species. In this study, we demonstrated that WGA generated smaller assembly and more fragmented contigs, probably due to the bias in the amplification process (Sabina et al., 2015). Albeit reduced, the quality of results remains high enough to allow certain types of studies, *e*.*g*. our nematode phylogenomic analysis. As a result, our study supports WGA as being a powerful tool in phylogenomic, population genomic and metagenomic studies of microscopic eukaryotes.

## Acknowledgements

X.Q. was funded by National Natural Science Foundation of China (32001876). Y.M.Z. was funded by the Oak Ridge Institute for Science and Education (ORISE) fellowship during part of this work. We thank Dr. Wilfrida Decraemer and Laboratory of Parasitology in Ghent University for providing nematode images, Dr. Kan Zhuo and Guokun Liu for providing nematode samples. We thank the Smithsonian Institution High Performance Cluster (https://doi.org/10.25572/SIHPC) for providing computational resources and support that have contributed to the research results reported in this publication.

## Supplementary material

Data available from the Dryad Digital Repository: https://doi.org/10.5061/dryad.b2rbnzsnt. The raw genomes available in NCBI with accession number PRJNA1128552.

## Notes

### Competing Interest Statement

The authors have declared no competing interest.

### Summary of Updates

New taxa have been added, phylogenetic trees redone. A more thorough review of the fossil calibration. Text rewritten to incorporate new results.

