## Supplemental Figures for "Phylogenomic Insights into the Evolution and Origin of Nematoda"

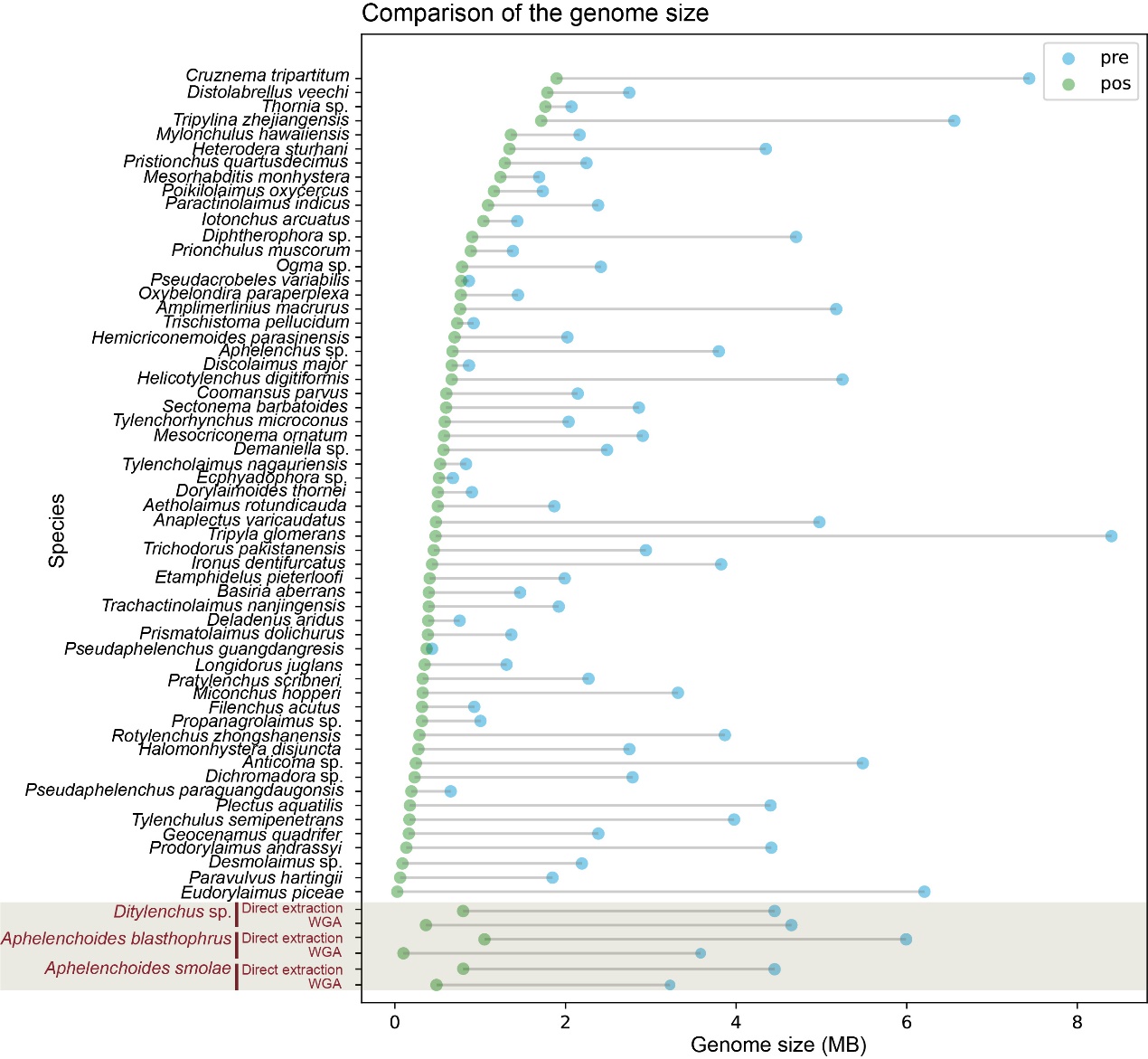

**Supplementary Figure 1.** The estimation of genome sizes for newly sequenced nematode species. Abbreviations: pre = the size of raw assembly, pos = the size after removing potential noise (fragments < 3k and coverage < 15). The three highlighted species were used to compare the assembly qualities between whole genome amplification (WGA) and direct DNA extraction methods.

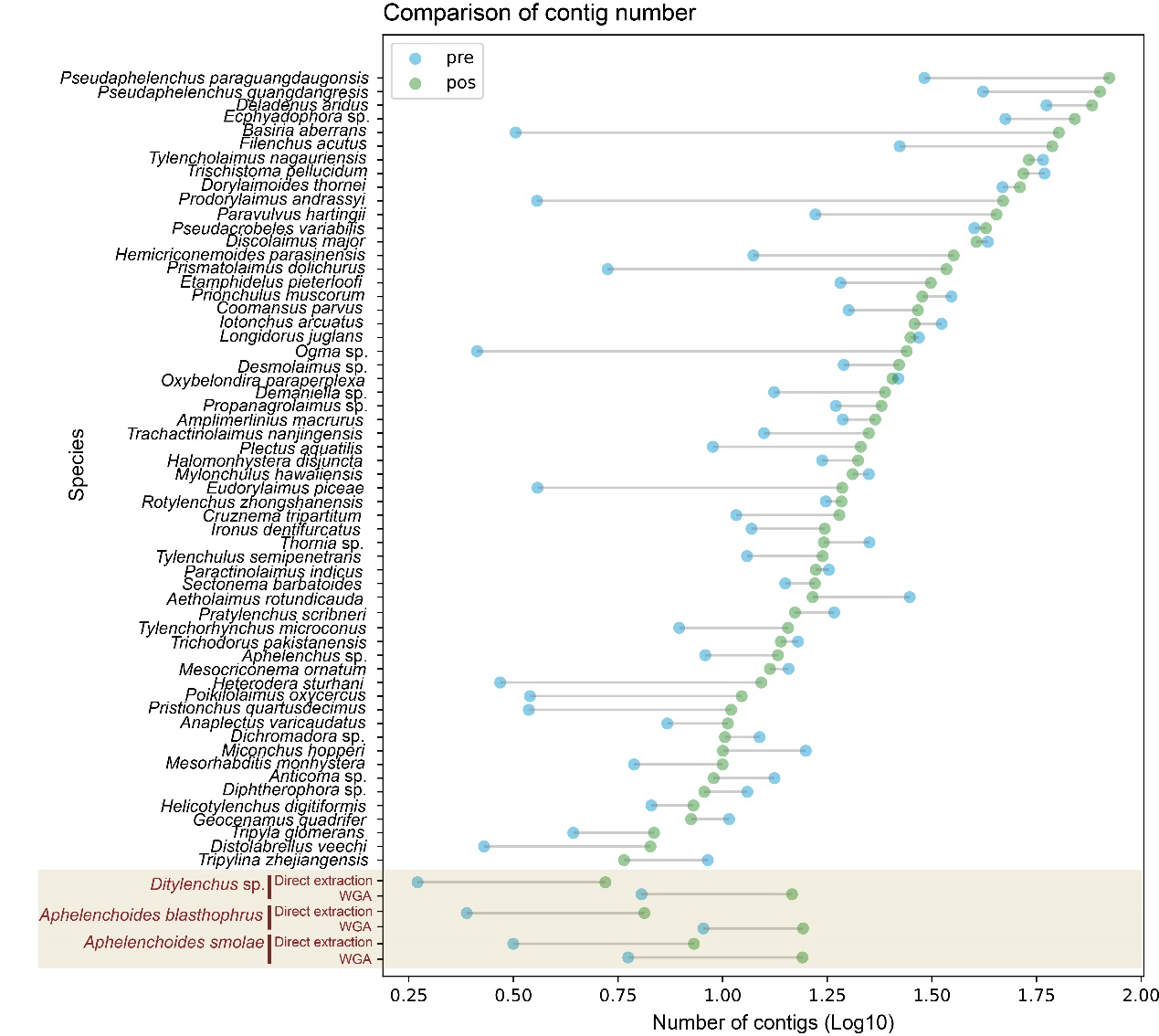

**Supplementary Figure 2.** The contig numbers of assemblies. Abbreviations: pre = the size of raw assembly, pos = the size after removing potential noise (fragments < 3k and coverage < 15). The three highlighted species were used to compare the assembly qualities between whole genome amplification (WGA) and direct DNA extraction methods. The longer line between pre and pos suggests higher proportion of short contigs in assembly.

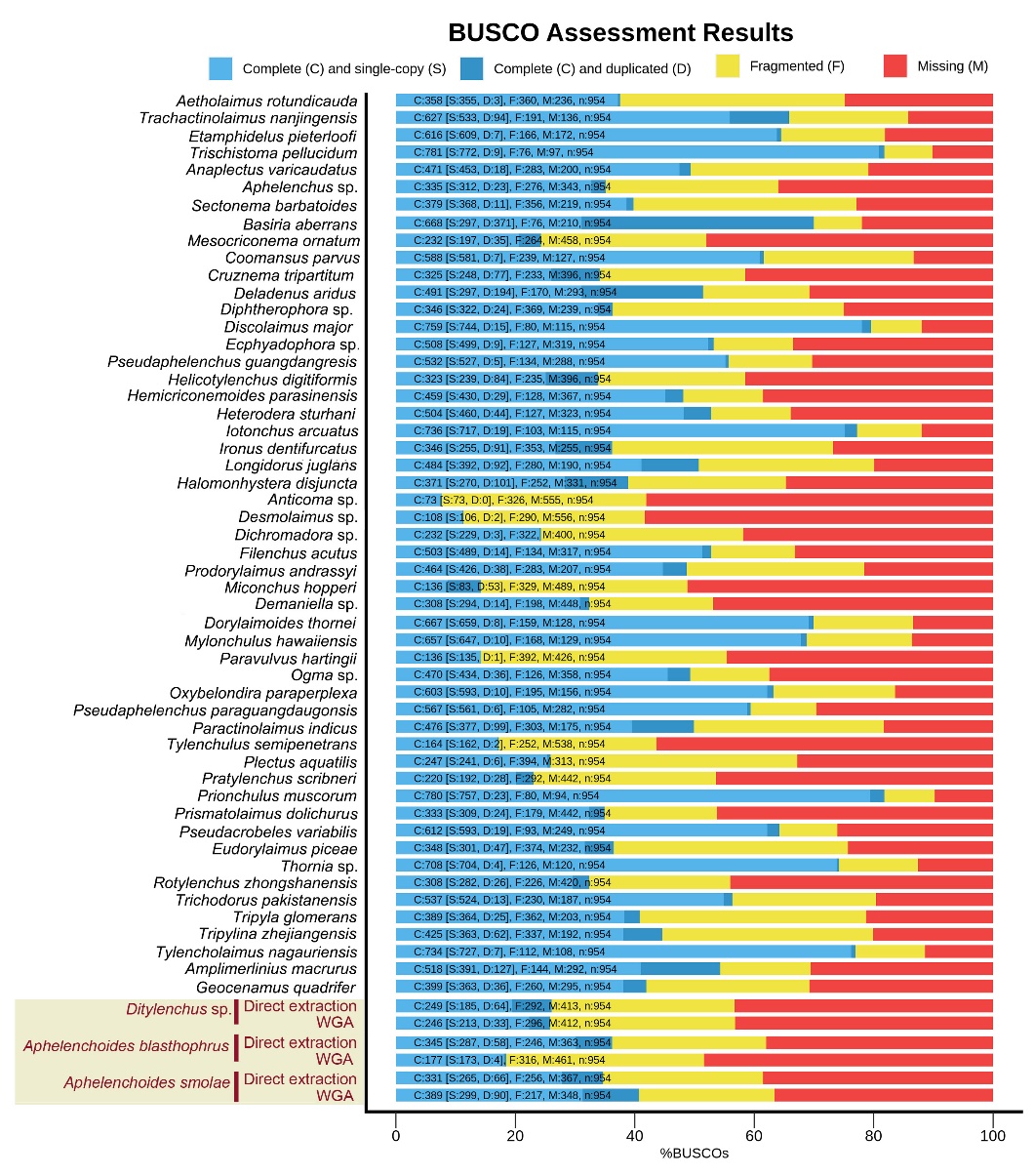

**Supplementary Figure 3.** The BUSCO completeness assessments for genomes (pre-trimming) obtained in this study using metazoa_odb10 database. The three highlighted species were used to compare the assembly qualities between whole genome amplification (WGA) and direct DNA extraction methods.

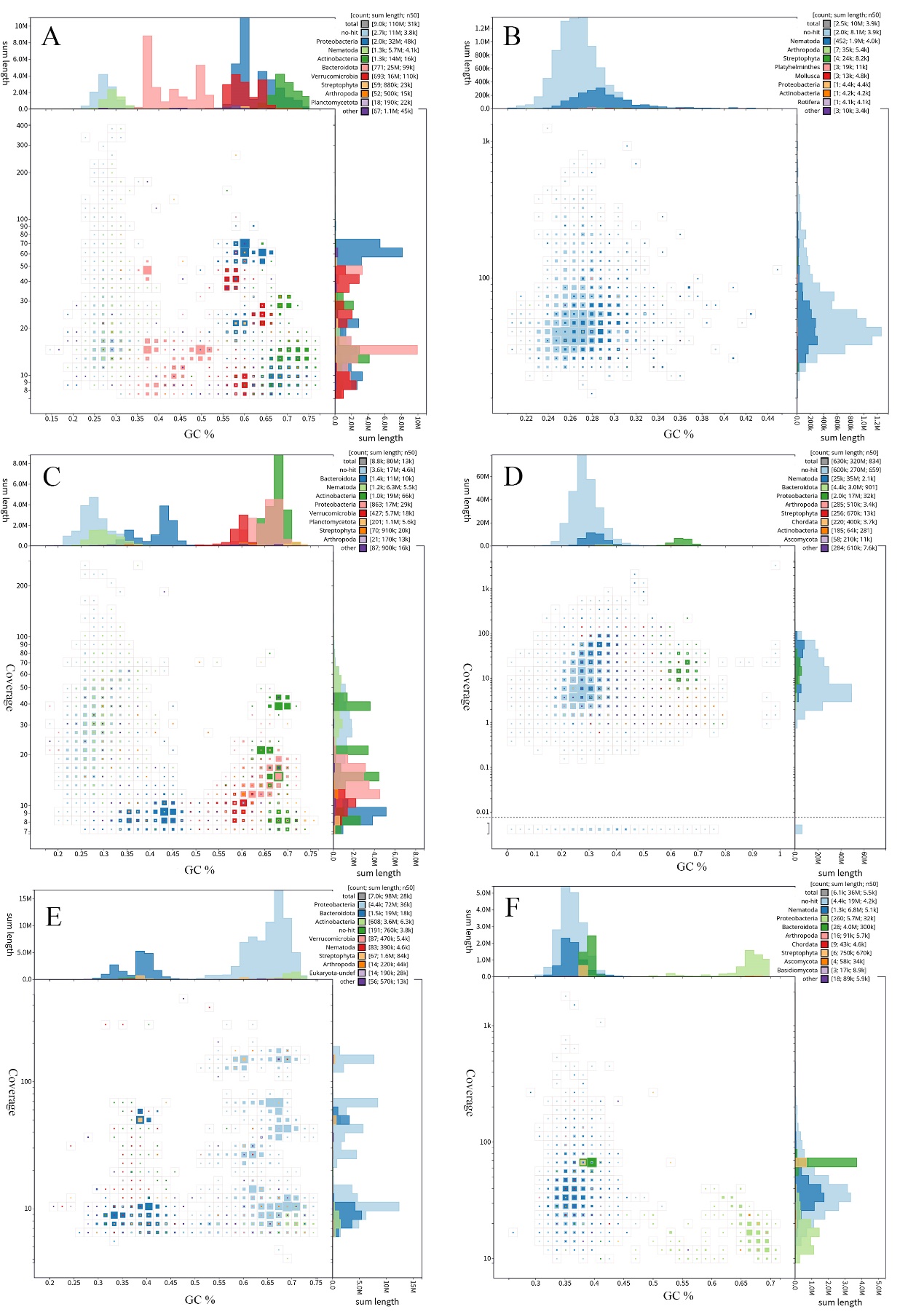

**Supplementary Figure 4.** The evaluation of sequencing contamination based on coverage and GC content of assembled contigs. The bacteria and Chordata contaminations are indicated with different colors, and the grey dots are unidentified contigs which are mostly nematodes. The more colored dots suggest more bacteria or Chordata contaminations. A, C, E: direct DNA extraction; B, D, F: whole genome amplification method. A, B: *Aphelenchoides blasthophrus*; C, D: *Aphelenchoides smolae*; E, F: *Ditylenchus* sp.

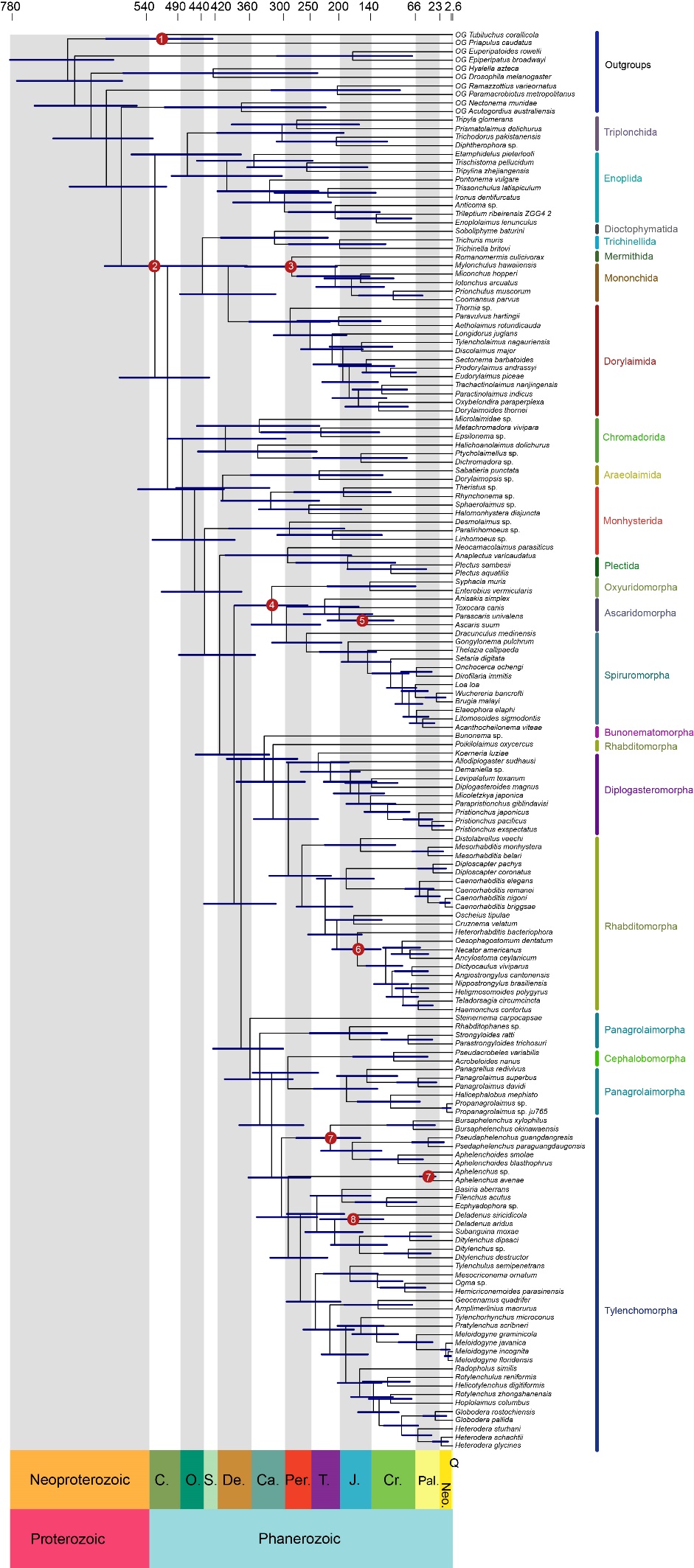

**Figure 5.** The Nematoda divergence time estimation performed by using MCMCTree on the PMSF_C60 topology. Node numbers and node bars represent 95% CIs of the estimated divergence times integrated from all MCMC runs. Number at nodes were constrained with the fossils records listed in supplementary Table S1. Abbreviation in geologic time scale: Q. = Quaternary, Neo. = Neogene, Pal.= Paleogene, Cr. = Cretaceous, J. = Jurassic, T. = Triassic, Per. = Permian, Ca. = Carboniferous, De. = Devonian, S. = Silurian, O. = Ordovician, C. = Cambrian.
